# A Ribosomal Marker-Based Metataxonomic Framework for Environmental Surveillance of Nematodes of Public Health Importance

**DOI:** 10.64898/2026.04.21.720024

**Authors:** Juan P. Zuluaga, Katherine Bedoya-Urrego, Juan F. Alzate

## Abstract

Metataxonomic analysis targeting the V4 region of the 18S rDNA gene, combined with molecular phylogenetic inference, was applied to detect nematode DNA of public health relevance in environmental matrices. A total of 25 mOTUs corresponding to six nematode taxa were detected in environmental samples from the Andean region of Colombia. Analysis of 12 water and sludge samples from wastewater treatment plants, 5 artisanal agricultural bioinputs, and 3 food samples revealed multiple species of public health significance: *Trichuris trichiura*, *Enterobius vermicularis*, *Ascaris* spp., and *Necator americanus.* We also confirmed zoonotic species, including *Angiostrongylus cantonensis* and *Trichinella* spp. These findings demonstrate that combining metataxonomics with molecular phylogeny provides a scalable molecular framework for the environmental surveillance of parasitic nematodes, overcoming the limitations of traditional morphological identification methods. This approach offers a replicable model for strengthening control and monitoring programs for parasitism in human populations.

## INTRODUCTION

Nematodes are ubiquitous metazoans found in terrestrial and aquatic habitats, parasitizing plants and animals including humans [1,2]. Approximately 30,000 species have been described, though total diversity may approach one million [3–5]. They exhibit remarkable trophic diversity, feeding on bacteria, fungi, algae, protozoa, and other nematodes, or living as facultative or obligate parasites [5,6]. In Colombia, nematode infections pose a significant public health problem, particularly among children [7], compounded by emerging anthelmintic resistance in both animal and human nematodes [8,9]. Intensive antiparasitic drug use has been selected for resistant organisms, compromising control programs [10], necessitating more sensitive molecular identification approaches.

Nematode species identification typically relies on morphological characteristics of adults, larvae, and eggs [11,12]. However, classical methods often produce nonspecific identifications, underestimating parasite circulation [6,12,13]. For instance, adult and larvae morphological similarities in mouthpart, esophagus and tail structures can obscure distinctions between species, and variations in body length may not adequately reflect species differences due to overlapping size ranges. Taxonomic resolution is frequently limited to genus level; for example, hookworm egg observation cannot differentiate *Necator americanus* from *Ancylostoma duodenale* [14]. Despite their ecological and physiological diversity, nematodes conserved morphology and small size provide few phylogenetically informative characters, many showing convergent evolution [6]. In environmental samples, clinically relevant nematode eggs may be misidentified as Acari (mite) eggs due to morphological similarities, potentially leading to diagnostic errors [15].

Conventional diagnostic methods cannot adequately detect parasitic nematode diversity [6]. In previous studies, metataxonomic approaches targeting ribosomal rDNA regions have demonstrated high sensitivity and specificity for cryptic or closely related species [16–18]. Though taxonomic resolution may be limited to genus level, metataxonomics offers reproducibility, scalability, automation, and cost-effectiveness [17], and facilitates parasite detection in complex environmental matrices [18]. While widely applied in prokaryotes [19–21], nematode metataxonomic studies remain emerging [3,22].

Accurate detection of parasitic nematodes in environmental matrices is critical for environmental surveillance and public health monitoring, yet remains constrained by limitations in sensitivity and taxonomic resolution. To address these challenges, this study implements metataxonomics coupled with phylogenetic analyses to detect nematodes of public health importance in environmental samples from the Andean region of Colombia.

We hypothesize that integrating metataxonomic profiling of the 18S rDNA V4 region with phylogenetic inference based on concatenated 18S–28S reference sequences enhances detection sensitivity and provides a versatile framework for identifying nematodes across heterogeneous environmental matrices. This approach enables robust taxonomic assignment and detection of morphologically cryptic and environmentally persistent taxa [17,22], while supporting the development of scalable molecular protocols for parasite surveillance within wastewater-based epidemiology and One Health frameworks.

## MATERIALS AND METHODS

### Sample Collection and DNA Extraction

Environmental samples were collected from four wastewater treatment plants (WWTPs), as well as from artisanal bioinputs and food sources. A total of 12 samples from WWTPs were obtained. The selected WWTPs are located in the Andean region, at elevations ranging from 1,080 meters above sea level (m a.s.l.) in Cali to 2,175 m a.s.l. [18]. The selected WWTPs collectively serve an estimated population of 5.5 million inhabitants. Two of these facilities San Fernando and Aguas Claras are situated in the Aburrá Valley metropolitan area and provide services to both the city and neighboring municipalities. The third plant, Cañaveralejo, is located in the city of Cali, while the fourth, El Retiro, is a smaller-scale facility serving a rural community in the municipality of El Retiro.

Three food samples from the Aburrá Valley and five organic bioinput samples from the cities of Cali and Medellín were processed using the same extraction protocol during 2023 and 2024, respectively. Prior to extraction, samples were homogenized using sterile instruments. DNA extraction was performed using the QIAGEN DNeasy PowerSoil Kit, processing 200 mg aliquots of environmental samples. The concentration and quality of purified DNA was determined by UV spectrophotometry, and samples were cryopreserved at −20 °C until use for PCR amplification [18].

### Metataxonomic Analysis

For DNA amplification, degenerate primers were used, designed to target the hypervariable V4 region of the eukaryotic 18S ribosomal gene (18S rDNA). The forward primer corresponded to the sequence 18S-V4Fw: (CCAGCAGCCGCGGTAATTCC) [23], the reverse primer used was 18S-V4Rev (RCYTTCGYYCTTGATTRA). These primers were successfully applied to the same sample set [18], in a study focused on protists. PCR amplification, genomic library preparation, and high-throughput sequencing services were outsourced to Macrogen Inc. (Seoul, South Korea), using the Illumina MiSeq platform configured for 300 bp paired-end reads.

Bioinformatic processing of the obtained sequences was performed using MOTHUR (v.1.44.3), following the protocol described by Rozo-Montoya et al. (2023). This process included the merging of paired-end reads, filtering of sequences containing ambiguous bases or shorter than 300 bp, removal of sequences with homopolymers longer than eight bases, clustering by sequence similarity, detection and removal of chimeric sequences, and construction of molecular operational taxonomic units (mOTUs) using a 97% similarity threshold.

Preliminary taxonomic assignment of the mOTUs was performed using the *classify.seqs* algorithm implemented in MOTHUR, with the SILVA (v.138) database serving as the reference [24]. Only mOTUs classified as eukaryotic were selected for subsequent BLASTn analysis [25], and phylogenetic studies. Sequencing quality indicators, including the number of high-quality sequences, mOTUs counts, and coverage estimators, were calculated using the *summary.single* command in MOTHUR. The raw amplicon sequences have been deposited in the NCBI SRA database under the Bioproject accession PRJNA976754.

### Species Selection and Bioinformatic Processing

For this analysis, nematodes of public health importance were selected, including species parasitic to humans, zoonotic species, [26,27]. In total, the reference database incorporated 49 species represented across 27 genera.

To obtain complete and high-quality rDNA sequences, nematode reference genomes of target species were downloaded from the NCBI database using its available datasets and a bioinformatic routine optimized for batch downloading. When no reference genome was available for a species of interest, complementary RNA-seq data were obtained from the NCBI Sequence Read Archive (SRA), followed by the extraction of contigs containing rDNA gene sequences using the Trinity program [28]. Neither genomic nor RNA-seq data were available for *Strongyloides ransomi*, instead partial 18S rDNA sequences (AB453327, OP288111) were included.

From the reference genomes, an annotation and extraction strategy was implemented to specifically retrieve the 18S and 28S ribosomal regions using Barrnap (v0.9) [29], combined with an auxiliary bioinformatic pipeline. The annotated and extracted sequences underwent quality screening to remove ambiguous, fragmented, or incomplete sequences. Only sequences with a combined length ≥ 1,400 bp across the 18S and 28S regions were retained. From the filtered set, a single consensus sequence with the highest overall quality and length was selected.

The resulting rDNA sequences were subjected to inspection and curation, including verification of orientation and length, format standardization, and consistent identifier assignment to ensure proper handling in subsequent analyses with SeqKit (v2.10.1) [30]. The 18S and 28S sequences were aligned separately using MAFFT (v7.215) [31], and the resulting alignments were inspected in AliView [32]. Finally, both alignment sets were concatenated using FASconCAT-G (v1.06.1) [33].

### Phylogenetic Analysis Using 18S and 28S rDNA Markers

To identify the putative nematode molecular operational taxonomic units (mOTUs) of interest in this study, a nucleotide identity–based search strategy was employed using BLASTn. For this purpose, the mOTUs generated with MOTHUR were compared against a local reference database containing concatenated 18S and 28S rDNA regions. mOTUs showing ≥95% sequence identity, representing the degree of match between aligned sequences, and a BLAST score ≥500, which reflects the overall quality and significance of the alignment according to the BLASTn algorithm, were considered valid candidates for phylogenetic analysis.

Reference sequences of 18S and 28S rDNA, together with the V4 region of the 18S rDNA from the putative mOTUs identified through BLASTn, were aligned using MAFFT (v7.215). The resulting alignment was manually inspected to identify and correct potential conflicting regions using AliView. Subsequently, maximum likelihood (ML) phylogenetic trees were constructed with the aligned sequences using IQ-TREE3 [34]. The robustness of branch support was evaluated using a dual approach: Ultrafast Bootstrap (UFBoot) and the Approximate Likelihood Ratio Test (aLRT), both computed with 5,000 replicates.

### Data Management and Presentation

Basic descriptive statistical analyses were performed using custom routines implemented in Python, including calculations of taxa presence–absence, occurrence frequencies, and genus- and species-level richness across samples and sampling sites. These summaries were used to support comparative interpretation of nematode diversity and distribution patterns.

Phylogenetic trees were visualized and edited using FigTree (v1.4.4) [35], where tree topologies were examined and clades of interest were selectively collapsed to enhance interpretability. Branch coloring and additional graphical refinements were subsequently applied using image editing software.

For graphical representation of spatial patterns, regional distribution maps were generated using QGIS (v3.40.11) [36]. Heatmaps summarizing taxa occurrence and relative abundance patterns were produced in R (v4.3.1) using the dplyr and ggplot2 packages.

## Results

### Construction of mOTUs and sequence processing

In each amplicon library, a minimum of 109,906 pairs of raw reads were obtained, with a maximum of 334,064 reads per sample. After merging and quality-filtering processes, the number of retained high-quality sequences ranged from 37,705 to 77,967. The number of molecular operational taxonomic units (mOTUs) detected across samples varied between 374 and 866. The coverage index ranged from 99.4% to 99.7%, indicating a high level of sampling of the expected theoretical diversity.

### Construction of a local database from Genomes

A total of 63 genomes representing 49 nematode species across 27 genera were analysed from the NCBI database. Genome sizes ranged from 42.5 to 656.4 Mb (median: 111.8 Mb).

Assembly continuity (N50) varied from 1.2 kb to 110.8 Mb (median: 1,095.2 kb), while fragmentation ranged from 2 to 167,310 contigs (median: 651). GC content ranged between 21.30% and 47.96% (mean: 36.68%). Most assemblies showed low levels of ambiguous bases (median: 0.33%), although higher values were observed in *Oesophagostomum dentatum*, *Romanomermis culicivorax*, and *Ancylostoma ceylanicum*.

The extraction of rDNA sequences using Barrnap was highly effective, achieving a success rate of 98.4% (62 out of 63 analyzed genomes). The only exception corresponded to *Enterobius vermicularis* (GCA_900576705), for which the software failed to identify ribosomal regions; thus, the dataset was complemented with nucleotide sequences obtained through partial RNA-seq annotation from assembly SRA_ERR310935 and Sanger sequences (FR687850, AF182295, JF934731, LC416069). A total of 14,875 rDNA sequences were recovered from genomes using this methodology: 6,276 corresponding to 18S and 8,599 to 28S. The concatenated consensus sequences (18S + 28S) ranged in length from 1,448 to 8,682 bp, with an average length of 4,925 bp, while the percentage of ambiguous nucleotides remained ≤ 0.03% in all cases.

### Taxonomic assignment of mOTUs by phylogenetic inference of rDNA

The initial analysis performed with MOTHUR generated a total of 16,045 mOTUs. Subsequently, the nucleotide identity search using BLASTn against the local reference database enabled the recovery of 933 hits that met the established filtering criteria (identity ≥ 95% and score ≥ 500). After removing redundant hits, 292 representative sequences were retained. These candidate sequences were aligned with MAFFT alongside the consensus reference sequences and subjected to phylogenetic inference. The general description of nematode genus per sample is described in Fig 1.

**Fig 1.**
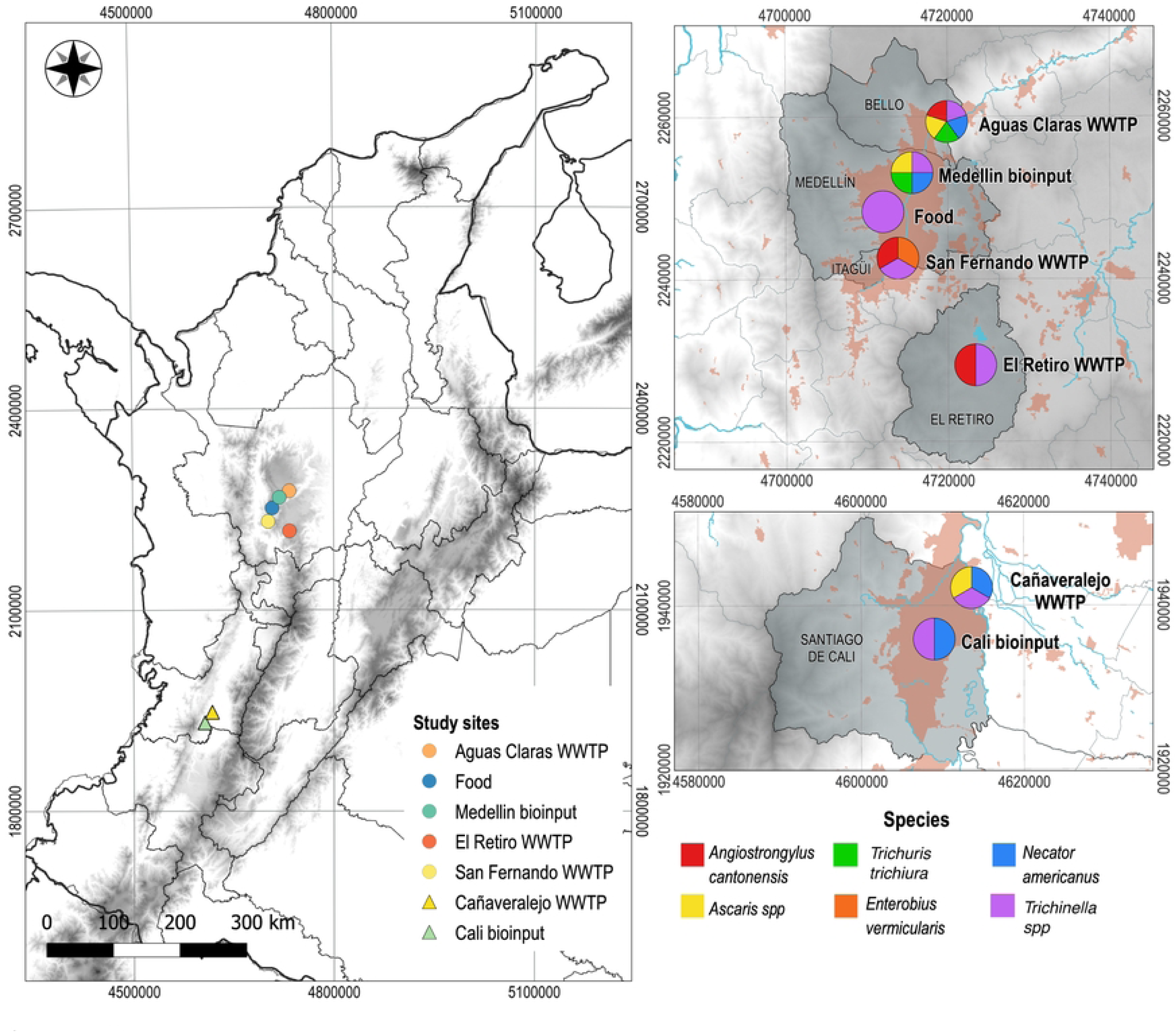
National and regional map of the Andean zone of Colombia showing all sampling locations included in the study. Coloured markers indicate the sites where nematode DNA was detected in wastewater treatment plants (WWTPs), biofertilizers, and food items collected between 2021 and 2024. This figure illustrates the geographic distribution of nematode presence, highlighting both urban and peri-urban areas. It provides a visual overview of the spatial heterogeneity of nematode contamination and identifies hotspots for potential epidemiological monitoring.

The maximum likelihood of phylogenetic analyses represented by the 25 mOTUs retained after filtering. The phylogenetic reconstruction of the nematodes of interest in this study showed high levels of statistical support, with UFBoot and aLRT values ≥ 95% for most clades, both calculated from 5,000 replicates. This degree of confidence allowed the consolidation of well-defined and clearly separated clades, consistent with previously reported phylogenetic structures described by other authors.[1,3,5,26]. Although it was necessary to supplement certain lineages not derived from reference genomes, the individual clade supports were sufficiently robust to allow a reliable assignment of the mOTUs included in the analysis. In Clade I, the presence of *T. trichiura* was confirmed with high phylogenetic support (UFBoot and aLRT ≥ 95%). For *Trichinella* spp. support values were slightly lower but still sufficient to enable reliable genus-level assignment. *For Trichinella spp*., phylogenetic analysis revealed clear separation from closely related species, with the mOTUs clustering within the clade comprising *T. britovi* and *T. pseudospiralis* (Fig 2).

**Fig 2.**
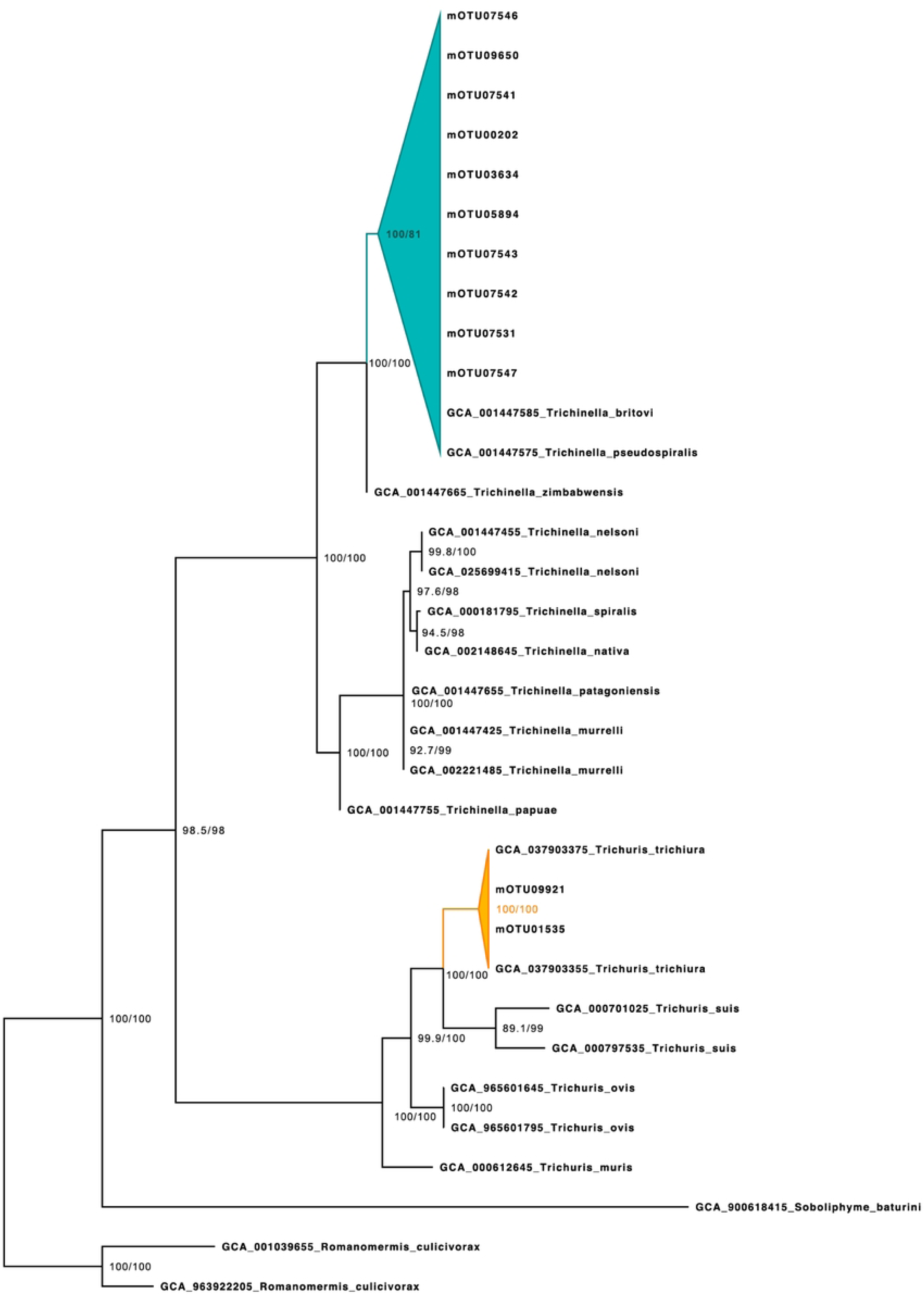
Phylogenetic analysis of Clade I for taxonomic assignment. Maximum-likelihood phylogenetic tree of Clade I, based on concatenated rDNA sequences (18S + 28S). Molecular operational taxonomic units (mOTUs) identified in the samples are labeled on the tree. High support values (UFBoot and aLRT ≥ 95 %) demonstrate the reliability of genus-level and, in some cases, species-level assignments. This clade includes nematodes of clinical and zoonotic relevance, supporting the utility of molecular monitoring in environmental matrices.

In Clade III, *E. vermicularis* was assigned with high confidence (UFBoot and aLRT ≥ 95%) and represented by five mOTUs. For the genus Ascaris, phylogenetic resolution did not allow differentiation between *A. suum* and *A. lumbricoides* due to their close evolutionary relationship (Fig 3).

**Fig 3.**
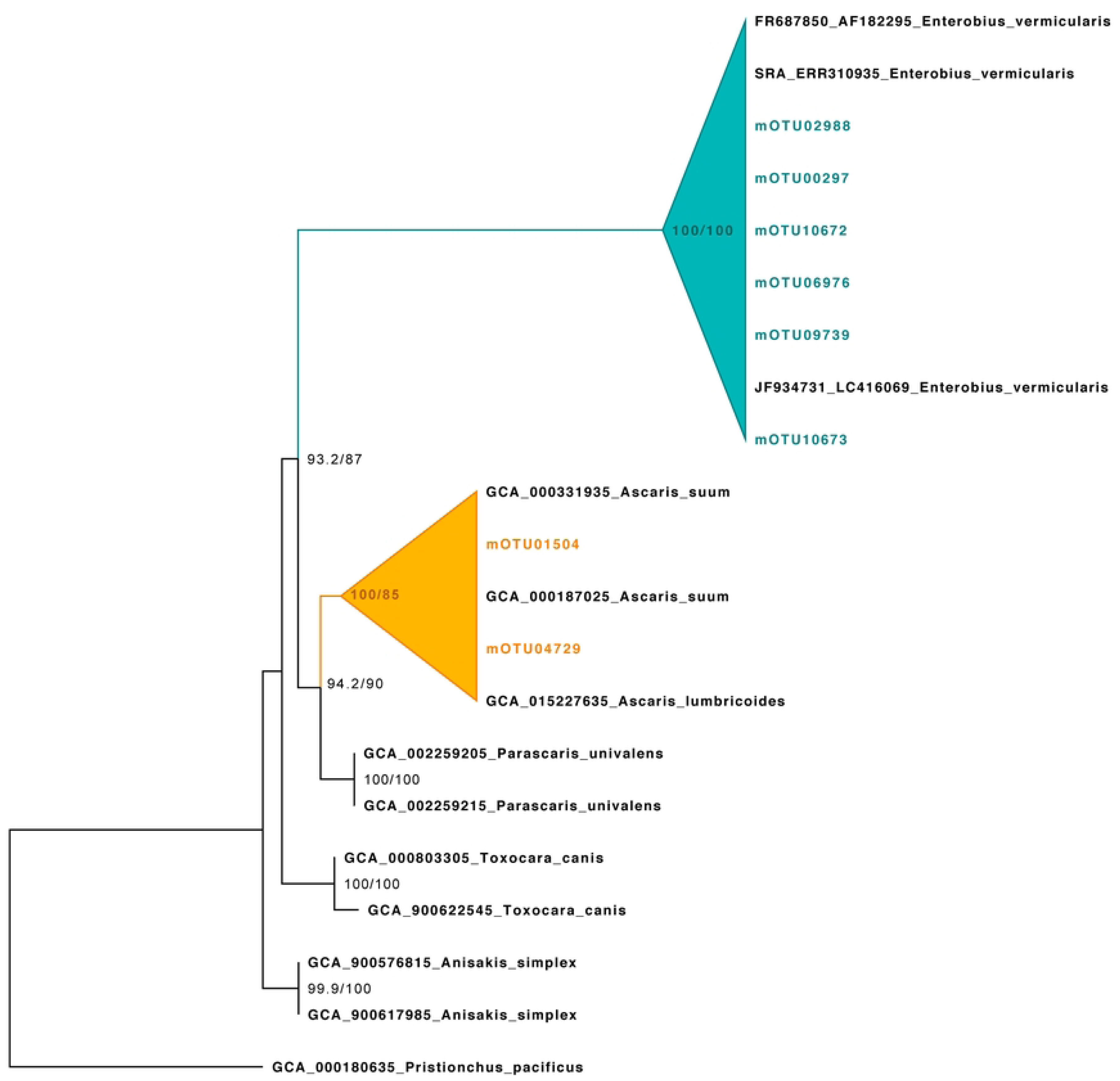
Phylogenetic analysis of Clade III for taxonomic assignment. Maximum-likelihood phylogenetic tree of Clade III, showing the placement of mOTUs identified in the study. The figure highlights intra-clade diversity and reveals phylogenetic relationships among nematodes present in wastewater and biofertilizer samples. Branch support values indicate the robustness of the assignments and provide confidence in interpreting potential transmission pathways.

In Clade IV, no species were confidently detected with support values above the significance threshold (UFBoot and aLRT ≥ 95%). Detailed results for this clade are therefore provided in the supplementary material (S1 Fig).

Finally, in Clade V, *A. cantonensis* were assigned with robust support (UFBoot and aLRT ≥ 95%). For *N. americanus*, support was slightly lower, although clustering within a well-defined clade allowed confirmation of its taxonomic identity (Fig 4).

**Fig 4.**
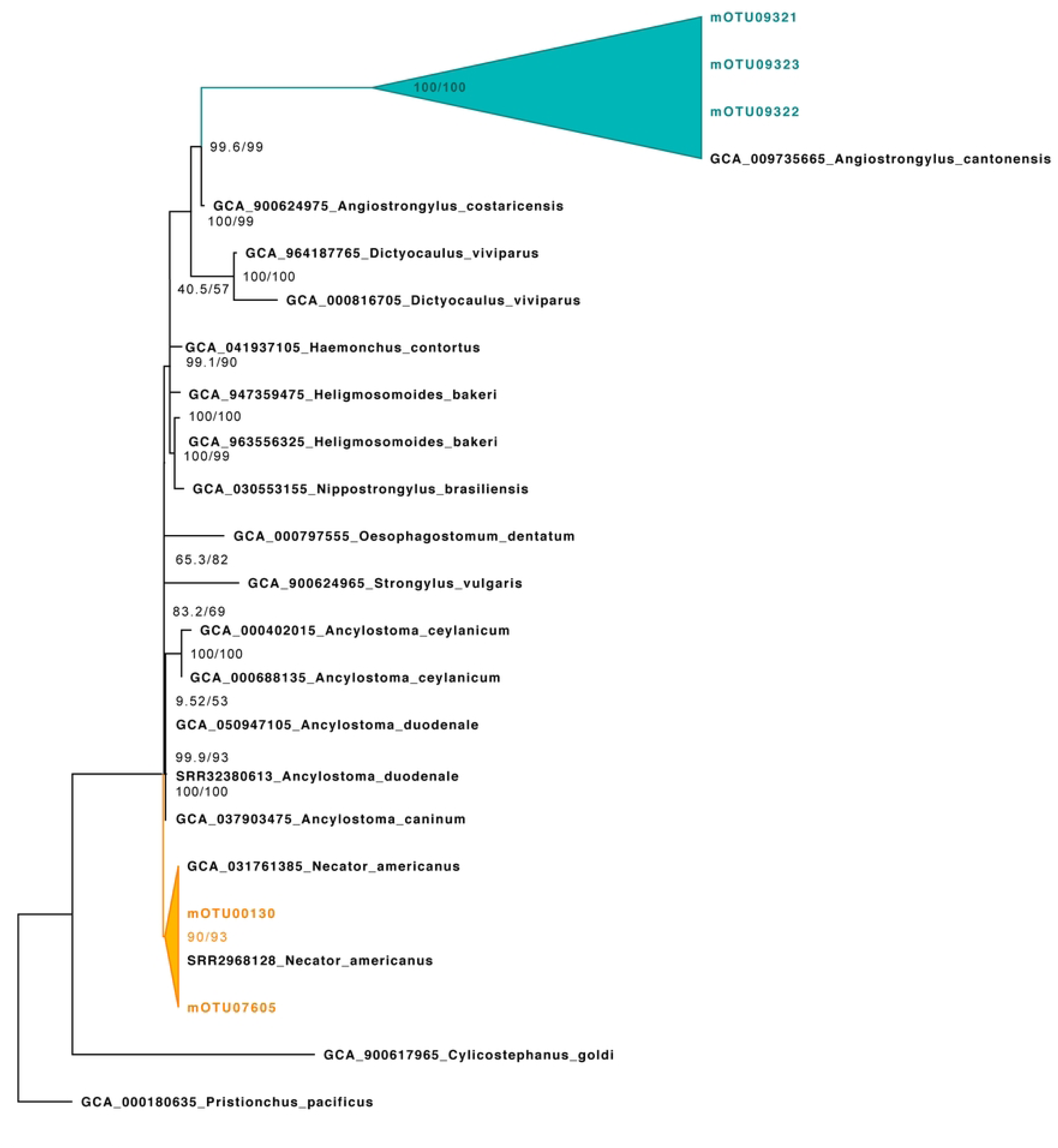
Phylogenetic analysis of Clade V for taxonomic assignment. Maximum-likelihood phylogenetic tree of Clade V, illustrating the placement of mOTUs detected in environmental samples. Support values (UFBoot and aLRT from 5,000 replicates) demonstrate reliable phylogenetic inference. This clade includes nematodes with varying degrees of public health significance, highlighting the importance of environmental surveillance for both human and zoonotic pathogens.

### Temporal Dynamics and Distribution of Nematode DNA in Wastewater Treatment Plants (WWTPs)

The occasional presence of nematode DNA was observed in the WWTPs, showing defined spatial distribution patterns at each study site (Fig 5). In Aguas Claras WWTP, a consistent pattern of nematode DNA detection over time was observed. In the influent water from 2021 (sample K7F4002), *Trichinella* spp. were detected. The biosolids from this plant appeared to act as a reservoir, with *N. americanus*, *Trichinella* spp., and *Ascaris* spp., detected in the same year (samples K7B2001 and K7B2002). In the 2023 influent (sample M9F4003), *Trichinella spp*., and *A. cantonensis* were identified. In contrast, the biosolids for the same period (sample M9B2003) contained *T. trichiura*, *Trichinella* spp., *N. americanus*, and *Ascaris* spp. (Fig 5, S2 Fig).

**Fig 5.**
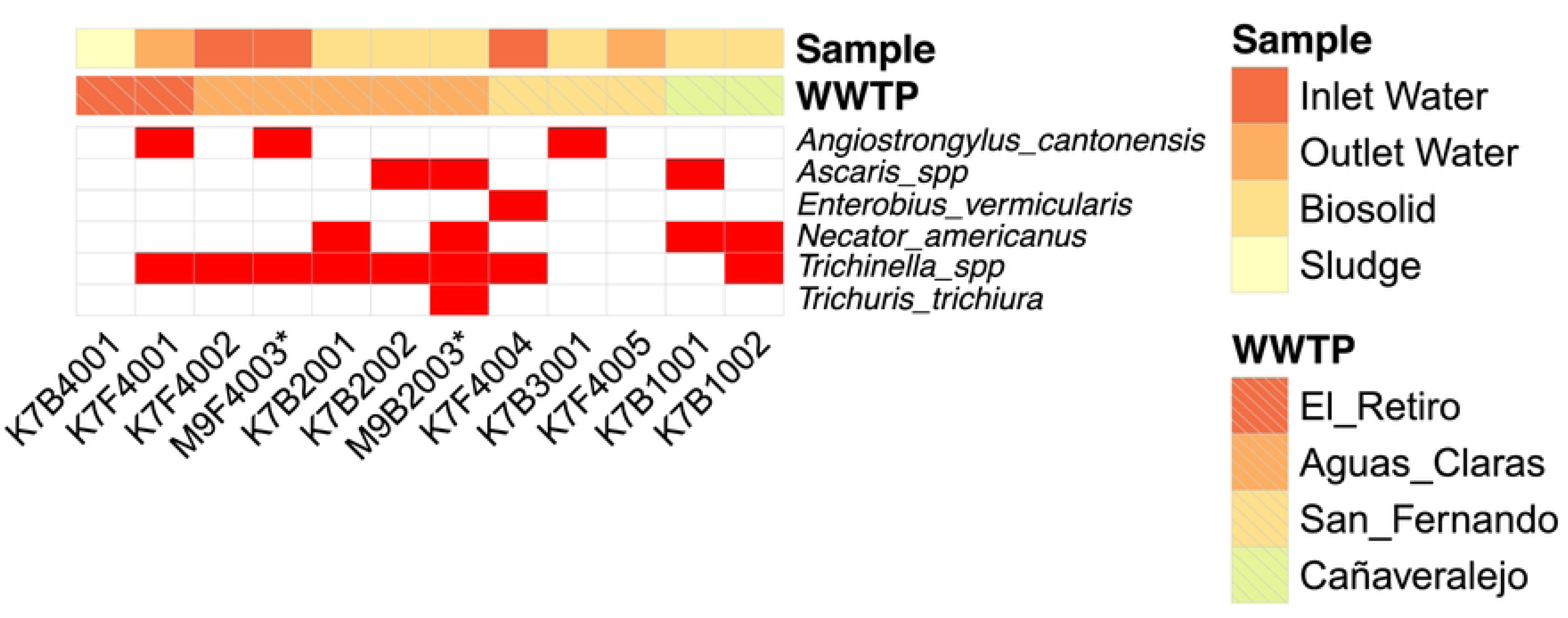
Comparative heatmaps of nematode DNA detection across wastewater treatment plants. Heatmaps illustrate the presence and absence of nematode DNA across all analyzed matrices between 2021 and 2024. Wastewater treatment plants (WWTPs), showing spatial and temporal patterns of detection and identifying facilities that act as persistent reservoirs of nematode genetic material (* indicates samples collected in 2023).

In San Fernando WWTP, during 2021, the influent water (sample K7F4004) contained *E. vermicularis*, and *Trichinella* spp. In the corresponding biosolid (sample K7B3001), A. cantonensis was detected, while the effluent showed no detectable nematode DNA (Fig 5, S3 Fig).

In Cañaveralejo WWTP (Cali), biosolid analysis revealed the persistence of *Ascaris* spp., *N. americanus*, in sample K7B1001, and the detection of nematode DNA corresponding to *N. americanus*, and *Trichinella* spp. in sample K7B1002 (Fig 5, S4 Fig).

Finally, in El Retiro WWTP, residual sludge (sample K7B4001) showed no detectable nematode DNA, whereas the effluent water (sample K7F4001) contained *Trichinella* spp., and *A. cantonensis*. (Fig 5, S5 Fig).

### Presence of Nematode DNA in Bioinputs and Foods

In the analysis of commercial bioinputs, DNA from species of public health relevance was detected in samples N8C1002, N8C1003, and N8C2001, all of which tested positive for *Trichinella* spp. Specifically, sample N8C1002 also exhibited a more diverse parasitic community, with the detection of nematode DNA corresponding to *N. americanus*, and *Ascaris* spp. In N8C1001, *N. americanus* was additionally detected, while in sample N8C2001, *N. americanus* was identified in both samples.

Regarding the food samples analyzed in 2023, only one sample (M9A2001) contained DNA corresponding to *Trichinella* spp. (Fig 6).

**Fig 6.**
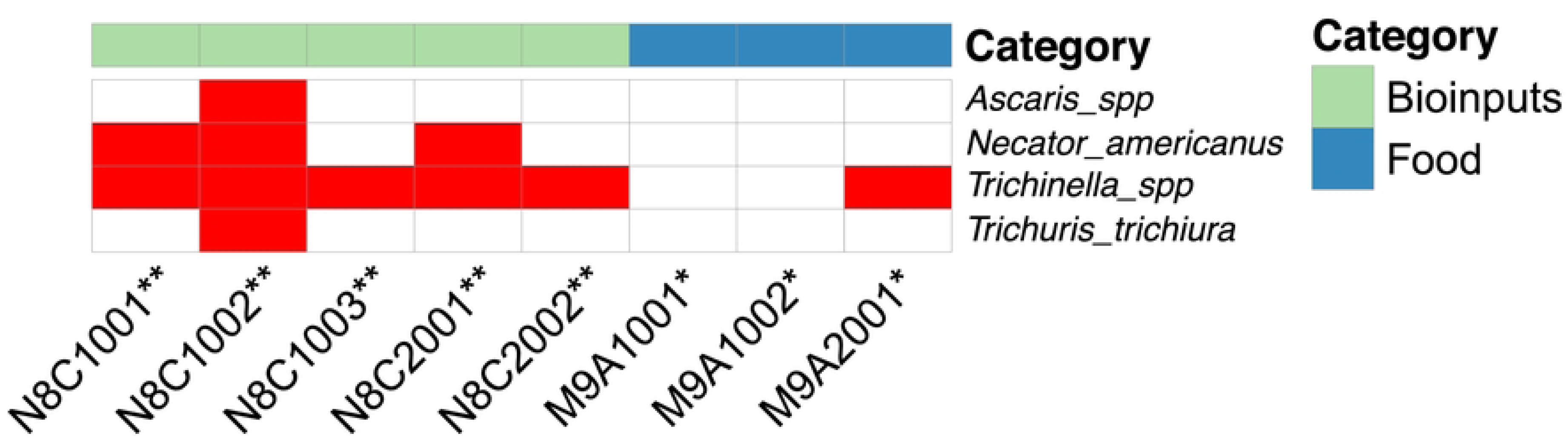
Comparative heatmaps of nematode DNA detection across biofertilizers, and food samples. Commercial biofertilizers and food items, evidencing potential parasite transmission routes through the reuse of treated or untreated biosolids in agriculture (* samples from 2023; ** from 2024).

## Discussion

### Wastewater-based epidemiological surveillance and relevance of nematodes as environmental markers

Wastewater-based epidemiology (WBE) has emerged as an effective approach for monitoring the circulation of infectious agents within human populations, allowing the detection of pathogens shed into urban sanitation systems and providing an indirect representation of community-level infection dynamics [18,20]. Traditionally applied to viruses, bacteria, and protozoa, WBE is increasingly recognized as a promising framework for monitoring helminths of public health importance. In this context, the application of metataxonomic approaches enables the detection of nematode DNA in complex environmental matrices such as wastewater and biosolids, providing a scalable strategy for environmental parasite surveillance.

In the present study, the detection of nematodes such as *Trichuris trichiura*, *Ascaris* spp., *Enterobius vermicularis*, and *Necator americanus* in wastewater treatment plants indicates that these systems act as environmental reservoirs of helminth DNA originating from human populations. The presence of these taxa is consistent with their known transmission routes, primarily associated with fecal contamination and inadequate sanitation [37]. The persistence of parasitic structures such as *Ascaris* and *Trichuris* eggs in wastewater systems has been widely documented due to their high environmental resistance and ability to accumulate in sludge and biosolids during treatment processes [38].

The taxa detected in this study also correspond broadly with national epidemiological records. According to the National Survey of Intestinal Parasitism in the School Population 2012–2014 (ENPI), the most frequently reported intestinal nematodes in Colombia are *Trichuris trichiura*, *Ascaris lumbricoides*, and hookworms (*Necator americanus* / *Ancylostoma duodenale*) [7]. The recurrent detection of *T. trichiura*, *Ascaris* spp., and *N. americanus* DNA in wastewater and biosolids therefore reflects the continued circulation of these geohelminths in the population and highlights their usefulness as indicators of fecal contamination and sanitation deficiencies. Notably, *A. duodenale* DNA was not detected in the analyzed samples, suggesting that in the studied regions the dominant hookworm component may correspond primarily to *N. americanus*.

Beyond these well-known soil-transmitted helminths, the detection of zoonotic taxa such as *Angiostrongylus cantonensis* and and *Trichinella* spp. expands the spectrum of parasites identifiable through environmental molecular monitoring. *A. cantonensis*, an emerging zoonotic parasite in Latin America associated with human angiostrongyliasis and eosinophilic meningoencephalitis [39,40], was detected in multiple matrices in this study. Its presence in wastewater suggests potential environmental circulation mediated by intermediate hosts or contamination pathways not captured through conventional clinical surveillance.

Taken together, these findings indicate that wastewater treatment systems function not only as sanitation infrastructures but also as environmental observatories for pathogen circulation. In Colombia, surveillance of soil-transmitted helminths remains largely focused on clinical reporting and periodic deworming campaigns, with limited integration of environmental data into public health monitoring systems. In this context, the metataxonomic detection of nematode DNA in wastewater provides an additional layer of information that could complement existing surveillance frameworks and support more proactive approaches to parasite monitoring within a One Health perspective.

### Bioinputs and food: parasitic risks in agricultural systems

The increasing commercialization of agricultural bioinputs derived from organic residues has expanded their use as fertilizers, soil conditioners, and microbial amendments in both small-scale and industrial agricultural systems. These products, often produced from treated organic matter or recycled waste streams, are widely promoted as sustainable alternatives to conventional agrochemicals. However, when sanitary controls are insufficient or production processes are poorly standardized, bioinputs may act as vehicles for the persistence and dissemination of microorganisms and parasitic DNA across agricultural environments. In this context, the evaluation of commercially available bioinputs and their potential role in the environmental circulation of parasites becomes particularly relevant, especially when these products are applied to soils used for food production [38,41].

Geohelminths, such as *Ascaris* spp., *Trichuris trichiura*, and *Necator americanus*, possess resistant structures that allow them to persist in environmental matrices such as water, soil, and biosolids. When these residues are used as bioinputs without proper treatment, the transmission cycle can be facilitated by contaminating agricultural soils and crops intended for human or animal consumption, thereby sustaining parasitic transmission cycles [38,41,42]. This dynamic perpetuates cross-contamination between the sanitation, agricultural, and food systems, creating an ecological circuit for parasitic persistence.

Although biological and thermal treatments significantly reduce microbial loads, various studies have demonstrated the residual presence of oocysts, cysts, and parasitic DNA even after conventional purification processes, highlighting their structural resistance [41]. Therefore, the production and use of bioinputs derived from treated sludge should include advanced sanitization processes and effectiveness controls, aimed at interrupting parasite transmission cycles and reducing associated risks [38,43].

In this regard, the safe management of bioinputs requires the integration of environmental, health, and food surveillance, ensuring that the reused residues do not pose a public health risk. Likewise, food represents a secondary exposure route, arising from the use of non-sanitized bioinputs for fertilization or from contact with contaminated environmental matrices, such as irrigation water. This connection underscores the need to assess parasitic traceability throughout the agri-food chain.

Accordingly, the molecular monitoring of nematodes and other parasites in reused matrices constitutes a key tool to strengthen agricultural biosafety, protect community health, and promote sustainability in wastewater treatment and reuse systems [42].

### Reference databases and limitations for species-level identification

The technical and technological feasibility of applying metataxonomic approaches to nematodes remains an emerging challenge. Although these methodologies are well established for bacteria and fungi, their development in parasites is still in its early stages. Nevertheless, the results obtained in this study demonstrate that it is possible to overcome, at least partially, the current limitations and move toward the standardization of specific protocols for the identification of nematodes in environmental matrices.

In this study, the combined use of the hypervariable V4 region of the 18S rDNA gene and concatenated 18S–28S reference sequences provided consistent taxonomic resolution and sufficient phylogenetic support to resolve several taxa at the genus level and, in some cases, at the species level [6,11].

The construction of the local reference database, derived from complete genomes and curated ribosomal sequences, was a key component of this study and simultaneously represented one of the main methodological challenges. Although the taxonomic coverage achieved was close to 100% of the nematodes included, the limited availability of complete genomes in public databases remains a major constraint for large-scale, high-resolution phylogenetic studies [5,22], and the biases associated with species diversity coverage will remain latent until this gap is resolved.

Through the strategy of annotating and extracting 18S and 28S ribosomal regions using Barrnap, more than 98% of the expected sequences were successfully recovered, generating 62 high-quality concatenated consensus sequences, which enabled the construction of a robust phylogenetic foundation. This advancement helps to address one of the most significant gaps in nematode metataxonomy: the lack of curated databases containing complete ribosomal information, which limits the taxonomic accuracy of inferences [22].

The maximum likelihood analysis allowed the identification of 25 mOTUs with strong statistical support (UFBoot and aLRT ≥ 95%), distributed across four clades and six nematode taxa. The topological congruence observed between the trees generated in this study and previously reported nematode phylogenies reinforces the robustness and reliability of the taxonomic assignments obtained [1,3,5,6]. These results support the reliability of the concatenated ribosomal marker system for resolving both deep evolutionary relationships and recent divergences [3,5].

The phylogenetic reconstruction confirmed the presence of six nematode taxa, several of which are of public health and zoonotic importance. Among the human intestinal nematodes, *T. trichiura*, *E. vermicularis*, and *N. americanus* were identified, all with high support values (≥ 95%).

Despite these advances, intraspecific resolution remains limited, especially in genera such as *Ascaris* and *Trichinella*, which exhibit low genetic divergence among closely related species. In these cases, taxonomic assignment was restricted to the genus level (spp.) due to the inability to distinguish between closely related species (*A. suum/A. lumbricoides*, *T. britovi/T. pseudospiralis*). Such limitations are consistent with previous metataxonomic studies of nematodes, where the resolving power of rDNA is considered moderate, yet sufficient for genus-level identification and effective for epidemiological surveillance [12,22].

This study demonstrates that ribosomal metataxonomics, when combined with phylogenetic validation using curated reference databases, constitutes a scalable framework for detecting parasitic nematode DNA across heterogeneous environmental matrices.

### Constraints and prospects

Despite the high sensitivity and reproducibility of the approach employed, intraspecific taxonomic resolution remains a challenge, particularly for lineages with low genetic divergence or highly conserved ribosomal sequences. Moreover, the detection of DNA does not necessarily imply the presence of viable or infectious organisms, so results should be interpreted in an ecological context, not solely at the molecular level.

The aim of this study was not to estimate parasite prevalence but to evaluate the feasibility of a metataxonomic framework for environmental surveillance across heterogeneous environmental matrices. Consequently, the results should be interpreted as evidence of methodological applicability rather than as a direct measure of parasite burden in the studied populations.

Future research should integrate higher-resolution markers, as well as phylogenomic strategies that allow more precise identification of cryptic and emerging species. Likewise, coupling spatial and temporal analyses could facilitate correlations between the presence of nematode DNA and local environmental or sanitary variables, thereby strengthening the predictive capacity of wastewater-based molecular surveillance.

## Supporting information

S1 Fig Phylogenetic analysis of Clade IV for taxonomic assignment. Maximum-likelihood phylogenetic tree of Clade I.

S2 Fig Comparative heatmaps of nematode DNA detection across wastewater treatment plants. Aguas Claras.

S3 Fig Comparative heatmaps of nematode DNA detection across wastewater treatment plants. San Fernando.

S4 Fig Comparative heatmaps of nematode DNA detection across wastewater treatment plants. Cañaveralejo

S5 Fig Comparative heatmaps of nematode DNA detection across wastewater treatment plants. El Retiro.

## Notes

### Competing Interest Statement

The authors have declared no competing interest.

## References

1. Koutsovoulos GD. Reconstructing the phylogenetic relationships of nematodes using draft genomes and transcriptomes [dissertation]. Edinburgh: University of Edinburgh; 2015. http://hdl.handle.net/1842/10558

2. Smythe AB, Holovachov O, Kocot KM. Improved phylogenomic sampling of free-living nematodes enhances resolution of higher-level nematode phylogeny. BMC Evol Biol. 2019;19(1):121. 10.1186/s12862-019-1444-x

3. Ahmed M, Roberts NG, Adediran F, Smythe AB, Kocot KM, Holovachov O. Phylogenomic analysis of the phylum Nematoda: conflicts and congruences with morphology, 18S rRNA, and mitogenomes. Front Ecol Evol. 2022;9:769565. 10.3389/fevo.2021.769565

4. de Ruiter PC, Neutel AM, Moore JC. Biodiversity in soil ecosystems: the role of energy flow and community stability. Appl Soil Ecol. 1998;10(3):217–28. 10.1016/s0929-1393(98)00121-8

5. van Megen H, van den Elsen S, Holterman M, Karssen G, Mooyman P, Bongers T, et al. A phylogenetic tree of nematodes based on about 1200 full-length small subunit ribosomal DNA sequences. Nematology. 2009;11(6):927–50. 10.1163/156854109x456862

6. Holterman MHM. Phylogenetic relationships within the phylum Nematoda as revealed by ribosomal DNA, and their biological implications [dissertation]. Wageningen: Wageningen University; 2007. https://edepot.wur.nl/121972

7. Ministerio de Salud y Protección Social, Universidad de Antioquia. Encuesta Nacional de Parasitismo Intestinal en Población Escolar Colombia, 2012–2014. Medellín: Facultad Nacional de Salud Pública, Universidad de Antioquia; 2015. https://www.minsalud.gov.co/sites/rid/Lists/BibliotecaDigital/RIDE/VS/PP/ET/encuesta-nacional-de-parasitismo-2012-2014.pdf

8. Geerts S, Gryseels B. Anthelmintic resistance in human helminths: a review. Trop Med Int Health. 2001;6(11):915–21. 10.1046/j.1365-3156.2001.00774.x

9. Matthews JB. Anthelmintic resistance in equine nematodes. Int J Parasitol Drugs Drug Resist. 2014;4(3):310–5. 10.1016/j.ijpddr.2014.10.003 PMID: 25516828

10. Vercruysse J, Albonico M, Behnke JM, Kotze AC, Prichard RK, McCarthy JS, et al. Is anthelmintic resistance a concern for the control of human soil-transmitted helminths? Int J Parasitol Drugs Drug Resist. 2011;1(1):14–27. 10.1016/j.ijpddr.2011.09.002 PMID: 24533260

11. Bogale M, Baniya A, DiGennaro P. Nematode identification techniques and recent advances. Plants. 2020;9(10):1260. 10.3390/plants9101260 PMID: 32992615

12. Roeber F, Jex AR, Gasser RB. Next-generation molecular-diagnostic tools for gastrointestinal nematodes of livestock, with an emphasis on small ruminants: a turning point? In: Rollinson D, editor. Advances in Parasitology. Vol 81. London: Academic Press; 2013. p. 267–333. 10.1016/B978-0-12-407705-8.00004-5

13. Powers T. Nematode molecular diagnostics: from bands to barcodes. Annu Rev Phytopathol. 2004;42:367–83. 10.1146/annurev.phyto.42.040803.140348 PMID: 15283671

14. Brooker S, Bethony J, Hotez PJ. Human hookworm infection in the 21st century. In: Rollinson D, editor. Advances in Parasitology. Vol 58. London: Academic Press; 2004. p. 197–288. 10.1016/S0065-308X(04)58004-1

15. Werneck JS, Carniato T, Andrade AG Jr, Tufik S, Andrade SS. Mites in clinical stool specimens: potential misidentification as helminth eggs. Trans R Soc Trop Med Hyg. 2007;101(11):1154–6. 10.1016/j.trstmh.2007.07.006 PMID: 17727906

16. Cabarcas F, Galvan-Diaz AL, Arias-Agudelo LM, García-Montoya GM, Daza JM, Alzate JF. Cryptosporidium hominis phylogenomic analysis reveals separate lineages with continental segregation. Front Genet. 2021;12:740940. 10.3389/fgene.2021.740940 PMID: 34712264

17. Garcia-Montoya GM, Galvan-Diaz AL, Alzate JF. Metataxomics reveals Blastocystis subtypes mixed infections in Colombian children. Infect Genet Evol. 2023;113:105478. 10.1016/j.meegid.2023.105478 PMID: 37406785

18. Rozo-Montoya N, Bedoya-Urrego K, Alzate JF. Monitoring potentially pathogenic protists in sewage sludge using metataxonomics. Food Waterborne Parasitol. 2023;33:e00210. 10.1016/j.fawpar.2023.e00210 PMID: 37808003

19. Bedoya K, Hoyos O, Zurek E, Cabarcas F, Alzate JF. Annual microbial community dynamics in a full-scale anaerobic sludge digester from a wastewater treatment plant in Colombia. Sci Total Environ. 2020;726:138479. 10.1016/j.scitotenv.2020.138479

20. Bedoya K, Coltell O, Cabarcas F, Alzate JF. Metagenomic assessment of the microbial community and methanogenic pathways in biosolids from a municipal wastewater treatment plant in Medellín, Colombia. Sci Total Environ. 2019;648:572–81. 10.1016/j.scitotenv.2018.08.119 PMID: 30121535

21. Porter TM, Hajibabaei M. Scaling up: a guide to high-throughput genomic approaches for biodiversity analysis. Mol Ecol. 2018;27(2):313–38. 10.1111/mec.14478 PMID: 29292539

22. Ahmed M, Back MA, Prior T, Karssen G, Lawson R, Adams I, et al. Metabarcoding of soil nematodes: the importance of taxonomic coverage and availability of reference sequences in choosing suitable marker(s). Metabarcoding and Metagenomics. 2019;3:e36408. 10.3897/mbmg.3.36408

23. Choi J, Park JS. Comparative analyses of the V4 and V9 regions of 18S rDNA for the extant eukaryotic community using the Illumina platform. Sci Rep. 2020;10(1):6451. 10.1038/s41598-020-63561-z PMID: 32300168

24. Quast C, Pruesse E, Yilmaz P, Gerken J, Schweer T, Yarza P, et al. The SILVA ribosomal RNA gene database project: improved data processing and web-based tools. Nucleic Acids Res. 2013;41(D1):D590–6. 10.1093/nar/gks1219 PMID: 23193283

25. Altschul SF, Gish W, Miller W, Myers EW, Lipman DJ. Basic local alignment search tool. J Mol Biol. 1990;215(3):403–10. 10.1016/S0022-2836(05)80360-2 PMID: 2231712

26. Coghlan A, Tyagi R, Cotton JA, Holroyd N, Rosa BA, Tsai IJ, et al. Comparative genomics of the major parasitic worms. Nat Genet. 2019;51(1):163–74. 10.1038/s41588-018-0262-1 PMID: 30397333

27. van Sluijs L, Geisen S, Lozano-Torres JL, Sterken MG, Vervoort MTW, Wilbers RHP. Unearthing roundworms: nematodes as determinants of human health. One Health. 2025;21:101103. 10.1016/j.onehlt.2025.101103

28. Grabherr MG, Haas BJ, Yassour M, Levin JZ, Thompson DA, Amit I, et al. Full-length transcriptome assembly from RNA-Seq data without a reference genome. Nat Biotechnol. 2011;29(7):644–52. 10.1038/nbt.1883 PMID: 21572440

29. Seemann T. Barrnap: bacterial ribosomal RNA predictor [software]. GitHub; 2013. https://github.com/tseemann/barrnap

30. Shen W, Le S, Li Y, Hu F. SeqKit: a cross-platform and ultrafast toolkit for FASTA/Q file manipulation. PLoS ONE. 2016;11(10):e0163962. 10.1371/journal.pone.0163962 PMID: 27706213

31. KaKatoh K, Standley DM. MAFFT multiple sequence alignment software version 7: improvements in performance and usability. Mol Biol Evol. 2013;30(4):772–80. 10.1093/molbev/mst010 PMID: 23329690

32. Larsson A. AliView: a fast and lightweight alignment viewer and editor for large datasets. Bioinformatics. 2014;30(22):3276–8. 10.1093/bioinformatics/btu531 PMID: 25095880

33. Kück P, Longo GC. FASconCAT-G: extensive functions for multiple sequence alignment preparations concerning phylogenetic studies. Front Zool. 2014;11(1):81. 10.1186/s12983-014-0081-x PMID: 25426157

34. Wong T, Ly-Trong N, Ren H, Baños H, Roger A, Susko E, et al. IQ-TREE 3: phylogenomic inference software using complex evolutionary models [preprint]. EcoEvoRxiv; 2025. 10.32942/X2P62N

35. Rambaut A. FigTree v1.4.4 [software]. Institute of Evolutionary Biology, University of Edinburgh; 2018. https://tree.bio.ed.ac.uk/software/figtree/

36. QGIS Development Team. QGIS geographic information system [software]. Beaverton: Open Source Geospatial Foundation; 2024. https://qgis.org

37. Loukas A, Maizels RM, Hotez PJ. The yin and yang of human soil-transmitted helminth infections. Int J Parasitol. 2021;51(13):1243–53. 10.1016/j.ijpara.2021.11.001 PMID: 34774540

38. Lu Q, He ZL, Stoffella PJ. Land application of biosolids in the USA: a review. Appl Environ Soil Sci. 2012;2012:201462. 10.1155/2012/201462

39. Bolaños F, Jurado-Zambrano LF, Luna-Tavera RL, Jiménez JM. Abdominal angiostrongyliasis, report of two cases and analysis of published reports from Colombia. Biomedica. 2020;40(2):233–42. 10.7705/biomedica.5043 PMID: 32673453

40. Gamarra-Rueda R, García R, Restrepo-Rodas DC, Pérez-García J. First identification of Angiostrongylus spp. in Lissachatina fulica and Cornu aspersum in Antioquia, Colombia. Biomedica. 2024;44(3):416–24. 10.7705/biomedica.7051

41. Amorós I, Moreno Y, Reyes M, Moreno-Mesonero L, Alonso JL. Prevalence of Cryptosporidium oocysts and Giardia cysts in raw and treated sewage sludges. Environ Technol. 2016;37(22):2898–904. 10.1080/09593330.2016.1168486 PMID: 27080207

42. Yakamercan E, Ari A, Aygün A. Land application of municipal sewage sludge: human health risk assessment of heavy metals. J Clean Prod. 2021;319:128568. 10.1016/j.jclepro.2021.128568

43. Laura F, Tamara A, Müller A, Hiroshan H, Christina D, Serena C. Selecting sustainable sewage sludge reuse options through a systematic assessment framework: methodology and case study in Latin America. J Clean Prod. 2020;242:118389. 10.1016/j.jclepro.2019.118389

